# Inversion of asymmetric histone deposition upon replication stress

**DOI:** 10.1101/2021.04.20.440573

**Authors:** Martijn R. H. Zwinderman, Thamar Jessurun Lobo, Petra E. van der Wouden, Diana C. J. Spierings, Marcel A. T. M. van Vugt, Peter M. Lansdorp, Victor Guryev, Frank J. Dekker

## Abstract

Following DNA replication, equal amounts of histones are distributed over sister chromatids by re-deposition of parental histones and deposition of newly synthesized histones. Molecular mechanisms balancing the allocation of new and old histones remain largely unknown. Here, we studied the genome-wide distribution of new histones relative to parental DNA template strands and replication initiation zones using double-click-seq. In control conditions, new histones were preferentially found on DNA replicated by the lagging strand machinery. Strikingly, replication stress induced by hydroxyurea or curaxin treatment, and inhibition of ATR or p53 inactivation, inverted the observed histone deposition bias to the strand replicated by the leading strand polymerase in line with previously reported effects on RPA occupancy. We propose that asymmetric deposition of newly synthesized histones onto sister chromatids reflects differences in the processivity of leading and lagging strand synthesis.

## Introduction

Posttranslational modifications (PTMs) of histones play important roles in regulation of nuclear organization and gene transcription (1). Histone PTMs are relatively stable and heritable during DNA replication (2, 3). Importantly, PTMs of parental histones differ from those of *de novo* synthesized histones (4). Therefore, near-symmetrical deposition of old and new histones during DNA replication is crucial to maintain similar chromatin states for the two sister chromatids following cell division. The functional importance of balanced histone inheritance was underscored by the observation that repressed chromatin domains are preserved by local re-deposition of parental histones (5). Clearly, mechanisms for accurate deposition of parental and new histones following DNA replication are crucial (6).

Recent studies have provided insight into histone deposition during DNA replication by using immunoprecipitation of PTMs on histones that are characteristic for either parental histones (H4K20me2) or new histones (H4K5ac) (7, 8). When this approach was combined with labeling of new DNA using the thymidine analogue 5-ethynyl-2’-deoxyuridine (EdU) to separate parental from newly synthesized DNA, a slight deposition bias of new histones to the lagging strand during DNA replication was revealed in mouse embryonic stem cells (7). A similar method was applied to budding yeast cells treated with hydroxyurea (HU), which revealed a slight bias of new histone deposition onto the leading strand (8). A clear explanation for these opposing findings has not been reported. Of note, both studies concluded that structural integrity of the replisome is essential to maintain near-symmetrical histone inheritance.

The structural integrity of the replication fork machinery is often compromised in cancer due to replication stress (9). Experimentally, replication stress can be induced by inhibition of the enzyme ribonucleotide reductase (RNR) with HU (10, 11). RNR inhibition results in a reduction of nucleotides required for DNA synthesis (12) and leads to uncoupling of the replicative helicase from the leading strand DNA polymerase (13). Helicase-polymerase uncoupling activates a DNA damage response (DDR), which involves the cell cycle checkpoint kinase ATR (14, 15) and p53-dependent transcriptional effects (16). However, the connection between replication stress and DDR pathways in relation to histone distribution following DNA replication is largely unexplored.

In this study, we have developed the double-click-seq protocol to enrich parental DNA bound to new histones. In contrast to previous methods involving immunoprecipitation of histone PTMs, this approach is based on metabolic labeling of new histones (Figure 1A). Double-click-seq provides stable labeling of *de novo* synthesized histones by co-translational incorporation of the methionine surrogate azidohomoalanine (AHA), which allows enrichment of nucleosomes that contain new histones following a click reaction with a biotin affinity tag. Using this methodology, we show that both replication stress as induced by HU and inhibition of the DDR invert the generally asymmetric distribution of new histones onto replicated DNA strands in human cells. We propose that asymmetric deposition of newly synthesized histones onto sister chromatids reflects differences in the processivity of leading and lagging strand synthesis.

**Figure 1.**
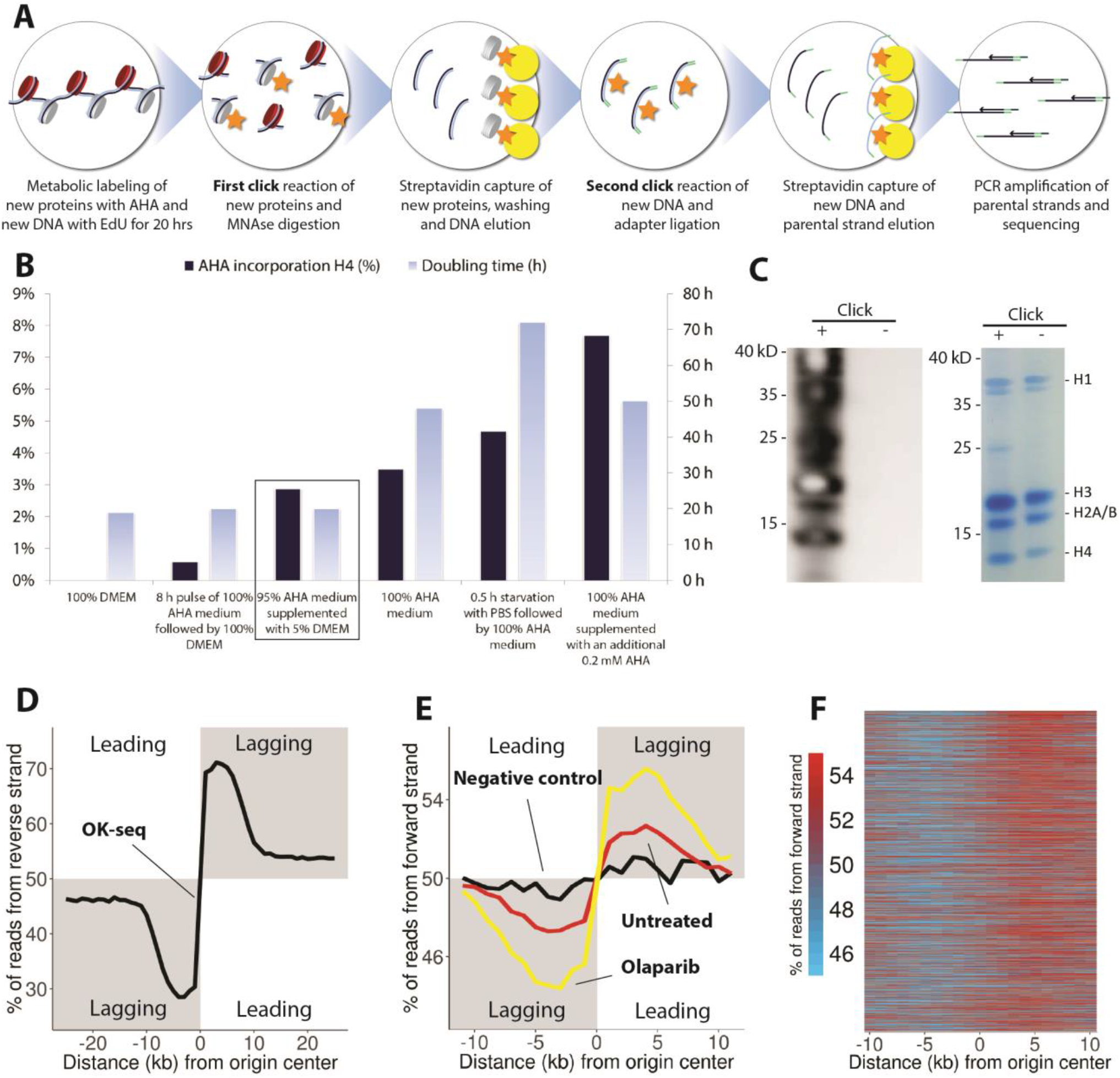
Double-click-seq reveals a bias in the deposition of new histones towards the DNA strand replicated by lagging strand machinery. (A) Schematic overview of the double-click-sequencing method. (B) Effect of different culture methods on cell doubling time and the extent of AHA incorporation in histone H4 determined by mass spectrometry. Culturing hRPE-1 in 95% AHA medium supplemented with 5% DMEM for 20 hours resulted in 3% AHA incorporation in histone H4 without a significant impact on cell doubling times (black box). (C) Western blot and gel electrophoresis analysis of labeled and unlabeled histones. Reactions were performed on nuclei with or without the click reaction as indicated above the blot and histones were extracted. Positions of histones are indicated to the right of the blot. (D) Average replication fork directionality at replication initiation zones. A plot including confidence intervals can be found in Supplementary Figure S1B. (E) Average bias of new histone deposition at replication initiation zones in the untreated condition (red line) and with olaparib treatment (yellow line). The black line shows the background signal in hRPE-1 cells (negative control, without second click reaction). Separate plots of individual replicates including confidence intervals can be found in Supplementary Figure S2. (F) Heatmap of new histone deposition bias at replication initiation zones in untreated condition.

## Materials and Methods

### Optimizing co-translational incorporation of azidohomoalanine (AHA)

#### Histone extraction

Nuclei were isolated as described for the Double-Click-seq experiment. Nuclei were resuspended in 400 μl 0.2 M H_2_SO_4_ and left at an end-over-end rotator for at least 30 minutes at room temperature. The samples were spun down in a tabletop centrifuge to remove nuclear debris at 16,000 g for 10 minutes. The supernatant (containing histones) were transferred to a fresh tube and while mixing 200 μl trichloroacetic acid (100%) was added drop by drop to the histone solution. The tube was inverted several times to mix the solutions and put on ice for 30 minutes. The histones were spun down in a tabletop centrifuge at 16,000 g for 10 minutes. The supernatant was carefully removed with a pipette and the histone pellet was washed with ice-cold acetone without disturbing it. The samples were centrifuged again at 16,000 g for 5 minutes and washing with acetone was repeated. The supernatant was carefully removed and the histone pellet was air dried for 20 minutes at room temperature. The pellet was then dissolved in an appropriate volume of water (typically 100 μl) and transferred to a fresh tube. The tube was centrifuged at 3400 g for 2 minutes and the supernatant was transferred to a new tube. The non-dissolved pellet contains mostly nonhistone proteins and other debris. The concentration was measured with the Bradford assay.

#### Propionic anhydride derivatization

Between 1 and 5 μg of the histone sample was aliquoted and diluted with 5 μL of 100 mM ammonium bicarbonate buffer (pH 8). The propionylation mixture was made by vortexing 75 μL of propionic anhydride with 25 μL 2-propanol. Next, 20 μL of this propionylation mixture was added to the histone sample, vortexed and centrifuged for a few seconds. The pH was checked with a pH indicator strip. If the pH was <8, ammonium hydroxide was added dropwise until the pH was approximately 8. Typically, adding 20 μL of ammonium hydroxide was a good starting point for the titration. The reaction was incubated at 51 °C for 20 minutes. The sample was concentrated to about 5 uL in a SpeedVac concentrator at room temperature for about 20 minutes. The sample was diluted with 5 uL of 100 mM ammonium bicarbonate buffer (pH 8) and derivatization was repeated. The reaction was then allowed to cool to room temperature without repeating the concentration step.

#### In solution trypsin digestion

Trypsin (Promega or NEB) was added to the histone sample at a 1:20 ratio (e.g., 5 mg of trypsin for 100 mg of histones) and the digestion was left at 37 °C for 6 hours. The digestion was stopped by adding 5 μL of glacial acetic acid to reach pH 3, which prevents trypsin from further digesting the histones. The sample was frozen at −80 °C to fully deactivate trypsin.

#### NanoChip LC–MS/MS QTOF

For the analysis of AHA incorporation of the histone peptides by LC–MS/MS, a quadruple time-of-flight mass spectrometer (QTOF, Agilent 6510) with a liquid chromatography-chip cube (# G4240) electrospray ionization interface was coupled to a nanoLC system (Agilent 1200) composed of a nanopump (# G2226A), a capillary loading pump (# G1376A) and a solvent degasser (# G1379B). Injections were performed with an autosampler (# G1389A) equipped with an injection loop of 40 μL and a thermostated cooler maintaining the samples in the autosampler at 4 °C during the analysis (# G1377A Micro WPS). The instrument was operated under the MassHunter Data Acquisition software (Agilent Technologies, Santa Clara, USA, version B.04.00, B4033.3). A chip (ProtID-Chip-150 II 300A, # G4240-62006) with a 40 nL trap column and a 75 μm × 150 mm analytical column filled with Zorbax 300SB-C18, 5 μm (Agilent Technologies, Santa Clara, USA) was used for peptide separation. The identification of peptides was based on data collected in auto MS/MS mode (2 GHz) using the following settings; fragmentor: 175 V, skimmer: 65 V, OCT 1 RF Vpp: 750 V, precursor ion selection: medium (4 m/z), mass range: 200–2500 m/z, acquisition rate for MS: 2 spectra/s, for MS/MS 3 spectra/s; MS/MS range: 50– 3000 m/z; ramped collision energy: slope 3.8, offset: 0, precursor setting: maximum 3 precursors/cycle; absolute threshold for peak selection was 1000; relative threshold was 0.01% of the most intense peak, active exclusion enabled after 1 selection, release of active exclusion after 0.6 min, precursors were sorted by abundance only. The MS/MS files were stored in centroid and profile mode. MS1 absolute threshold 50 and MS2 absolute threshold 35 were C646 49 2 2 applied to account for detector noise. A static exclusion range of 200–350 m/z for precursor selection was applied. Gas temperature (nitrogen) was 325 °C and gas flow was 5 L/min. The quantification of peptides was based on data collected in MS mode using the same settings except of the mass range 200–3000 m/z; acquisition rate 1 spectra/s. In both cases lock masses 1221.990 m/z (HP-1221; Agilent article number G1982-85001) and 299.294 m/z (Methyl Stearate; Agilent article number G1982-85003) were used to recalibrate spectra during the acquisition.

Tandem mass spectra were extracted, charge state deconvoluted and deisotoped by the MassHunter Qualitative Analysis software version B.05.00 (Agilent Technologies) and saved as mgf files. All MS/ MS data were analyzed using Phenyx (GeneBio, Geneva, Switzerland); version CYCLONE (2010.12.01.1). The fragment ion mass tolerance of 0.30 Da and a parent ion tolerance of 400 ppm were selected for a database search. Oxidation (+15.99) and replacement by AHA (−4.98) of methionine, acetylation (+42.01), methylation (+14.01), dimethylation (+28.03), trimethylation (+42.04), propionylation (+56.03) of lysine, methylation (+14.01), dimethylation (+28.03) of arginine and amidation of aspartic acid (−0.98) were specified in Phenyx as variable modifications. The ratio between AHA labeled and unlabeled peptides of histone H4 (79–92) KTVTAMDVVYALKR was calculated using the areas under the curve in the extracted ion chromatograph to give an estimate of AHA incorporation in new histones. Total histone H4 was calculated by summing the areas of AHA labeled and unlabeled H4 peptides.

### Double-click-seq

#### Incubation and harvest of cells

The experiments for all treatments were done in duplicate. As a result libraries from two biological replicates were generated for each condition, except p53Null+HU (material lost during library prep stage) and single-click negative control (sample requested during laboratory lockdown). Four T175 flask were each seeded with approximately 3 ⨯ 10^6^ cells per experiment. After overnight incubation, culture medium was removed and replaced by pre-mixed medium containing 0.95 × volume 100% AHA medium (DMEM, high glucose, no glutamine, no methionine, no cysteine (Gibco) was supplemented with 10% fetal bovine serum, 1% penicillin-streptomycin (10,000 units penicillin and 10 mg streptomycin/mL, Gibco), 4 mM GlutaMax™ (Gibco), 0.2 mM L-Cystine 2HCl, 1 mM sodium pyruvate, 0.2 mM AHA and 20 µM EdU) and 0.05 × volume of DMEM (see recipe above). For induction of replication stress either 1 µM olaparib, 100 nM curaxin (CBL0137), 150 or 200 µM hydroxyurea (HU), 0.5 µM ATR inhibitor (VE-821) or a combination was added to the culture medium. Cells were incubated for 24 h, after which the culture medium was removed and cells were washed twice with ice-cold phosphate-buffered saline (PBS). Cells were harvested with Trypsin-EDTA (Gibco) for 5 minutes at 37 °C. Trypsinization was quenched by the addition of medium and cells were collected by centrifugation at 200 g for 5 minutes. The pellet was resuspended in ice-cold PBS and centrifuged again at 200 g for 5 minutes. The cell pellet was stored at −80 °C until further use.

#### Nuclei isolation

The cell pellet was resuspended in 1 mL of ice-cold PBS containing 0.1% NP-40 supplemented with proteinase inhibitor cocktail (PIC) (cOmplete™, Mini, EDTA-free, Roche). The cell suspension was vortexed for five seconds at maximum speed. The lysed cell suspension was centrifuged for 10 seconds at 9000 g. The supernatant was discarded and the nuclei were resuspended in 1 mL of ice-cold PBS containing 0.1% NP-40 supplemented with PIC. The washed nuclei were centrifuged for 10 seconds at 9000 g and the supernatant was discarded.

#### First click reaction

The nuclei were resuspended in 250 µL nuclei buffer (320 mM sucrose, 5 mM MgCl_2_, 10 mM HEPES at pH 7.4) supplemented with PIC and 10 µL 100 mM (in methanol) biotin-PEG4-alkyne (Sigma-Aldrich) was added. Next, a pre-mixed solution of 5 µL 50 mM CuSO_4_ with 5 µL 250 mM THPTA (Sigma-Aldrich) was added, followed by 10 µL of a freshly prepared solution of 100 mM sodium ascorbate (17). The nuclei were placed in an end-over-end rotator at room temperature for 30 minutes. After that, the click reaction was repeated as described above using fresh reagents. Nuclei were pelleted for 10 seconds at 9000 g (to remove excess biotin, which would interfere with streptavidin bead pull-down later on) and the supernatant was discarded.

#### MNAse digestion

Nuclei were resuspended in 100 µL of MNAse buffer (50 mM Tris.HCl, 5 mM CaCl_2_, pH 7.9, supplemented with 100 µg/mL BSA) and warmed to 37 °C in a water bath. Then, 60 U micrococcal nuclease (MNase, New England Biolabs, 1U/µL in MNAse buffer) was added and cells were incubated at 37 °C for 5 minutes. MNase reactions were stopped by adding EDTA to a final concentration of 5 mM and by cooling on ice. Next, 100 µL PBS supplemented with 0.1% BSA and 0.01% Tween was added and the nuclei were pelleted by centrifugation for 10 seconds at 9000 g.

#### Streptavidin enrichment of labeled nucleosomes

The supernatant (containing the nucleosomes) was added to 30 µL streptavidin magnetic beads (Dynabeads™ MyOne™ Streptavidin C1, Thermo Scientific, pre-washed three times with PBS according to the instruction manual) and incubated at room temperature for 1 hour in an end-over-end rotator. The supernatant was removed by placing it on a magnet for 3-4 minutes. The beads were washed five times by resuspension in 200 µL PBS supplemented with 0.1% BSA and 0.01% Tween and placing them back on the magnet. After washing, the DNA bound to the captured nucleosomes was released. This was done by resuspending the beads in 100 µL PBS supplemented with 1 mg/mL proteinase K and incubating the suspension for 30 minutes at 50 °C (pipet up and down after 15 minutes to ensure a homogenous mixture). Then, 100 µL 5 M NaCl was added and the mixture was left rotating for 15 minutes at room temperature. Finally, 360 µL (1.8 ⨯ volume of sample) Ampure beads (Beckman) was added directly to the bead suspension to purify the released DNA according to the protocol of the manufacturer. The DNA was eluted in 80 µL water.

#### Second click reaction

1 µL 100 mM (in methanol) biotin-PEG4-azide was added to the eluted DNA, followed by a pre-mixed solution of 5 µL 50 mM CuSO_4_ with 5 µL 250 mM THPTA and 10 µL of a freshly prepared solution of 100 mM sodium ascorbate. The sample was placed in an end-over-end rotator at room temperature for 45 minutes. To discard DNA fragments larger than approximately 200 bp 70 µL (0.7 ⨯ volume of sample) Ampure beads were added and after 5 minutes incubation at room temperature the beads were discarded and the supernatant was transferred to a new tube. Then 180 µL (1.8 ⨯ original sample volume) Ampure beads were added and purification was completed according to the manufacturer’s protocol. The DNA was eluted in 55.5 µL water.

#### End prep and adaptor ligation

End repair, 5’ phosphorylation, dA-tailing and adaptor ligation was performed using the NEBNext® Ultra™ DNA library prep kit for Illumina®. After adaptor ligation the sample was added to 25 µL streptavidin magnetic beads (Dynabeads™ MyOne™ Streptavidin T1, Thermo Scientific) pre-washed three times with B&W buffer (5 mM Tris HCl pH 7.5, 0.5 mM EDTA, 1 M NaCl) according to the instruction manual).

#### Isolation of replicated double stranded DNA fragments

After 15 minutes incubation at room temperature in an end-over-end rotator, beads were washed four times with 200 μl 1x B&WT buffer (5 mM Tris HCl pH 7.5, 0.5 mM EDTA, 1 M NaCl, 0.05 % Tween 20) and once with 2 × B&WT buffer.

#### Isolation of replicated parental strands

The parental strands were then eluted from the beads by incubation with 100 μl alkaline buffer (0.1 M NaOH, 0,05 % Tween 20) for 1 minute (7). Elution of the parental strand was repeated twice for a total of three times and the pH of the combined alkaline supernatants was neutralized by adding acetic acid to a final concentration of 0.1 M and 2 mM EDTA pH 8.0. The sample was diluted to 1 mL with water and concentrated to approximately 20 uL with a Vivaspin® 2 5K centrifugal concentrator according to the manufacturer’s instruction (15 minutes centrifuging at 4000 g in a swing bucket). The DNA was recovered by performing a reverse spin at 3000 g for 2 minutes.

#### Library amplification and sequencing

Libraries were amplified using the NEBNext® Ultra™ DNA library prep kit for Illumina® using NEBNext® multiplex oligos for Illumina® (index primer set 1 or 2) with 13 PCR cycles. Purification was done as described in the manual (with 0.9 × Ampure beads) and eluted in 30 μl water. Libraries were quantified with Qubit™ dsDNA high sensitivity assay kit (Thermo) and further analyzed with the high sensitivity D1000 ScreenTape system (Agilent). Sequencing was performed on a NextSeq 500 with onboard clustering using a 75-cycle high output v2.5 kit (Illumina) to generate paired-end 41-bp-long reads.

#### Generating negative control and control of streptavidin enrichment of labeled nucleosomes

The washing fractions of the streptavidin enrichment step of labeled nucleosomes were kept and analyzed with gel electrophoresis (see Figure S1E). Only non-specific protein elution from the beads was observed, indicating that the stringent washing steps were sufficient to primarily enrich nucleosomes over other DNA-protein complexes, as others have observed as well (17). To test for absence of background signal the double-click-seq protocol was followed until streptavidin enrichment of labeled nucleosomes. After DNA elution of the beads, and an Ampure bead clean-up, the resulting double-stranded DNA was eluted in 55.5 µL water. This was followed by end prep, adaptor ligation and library amplification according to the NEBNext® Ultra™ DNA library prep kit manual. Sequencing, paired-end, 150 bp, was performed using an llumina NovaSeq 6000 device. Furthermore, performing double-click-seq with cells incubated with EdU but without AHA or *vice versa* did not result in DNA amplification.

### Inactivation of TP53 in hTERT-RPE-1 cells

hTERT immortalized human retinal pigmented epithelial cells (hTERT RPE-1, ATCC ®, CRL-4000™) were used. To mutate *TP53*, a single guide RNA (sgRNA) targeting exon 4 of *TP53* (5’-CTGTCATCTTCTGTCCCTTC-3’) was cloned into pSpCas9(BB)-2A-GFP (pX458, plasmid #48138, Addgene). pSpCas9(BB)-2A-GFP was a kind gift from dr. Feng Zhang (18). RPE-1 cells were transfected with pX458 using Fugene HD transfection reagent (Promega) according to the manufacturer’s protocol and 48 hours later cells were selected for 3 weeks with 10 mM Nutlin-3 (Selleck Chemicals). *TP53* was mutated in exon 4 by a 7 basepair deletion (TCA-TCT-T), leading to a frameshift at codon 97. *TP53* mutations in exon 4 were confirmed by Sanger sequencing.

### Quantification and Statistical Analysis

Scripts in this section are available at https://github.com/thamarlobo/histone_deposition_analysis.git

#### Raw data processing of OK-seq data

Quality trimming of raw OK-seq reads for RPE-1 hTERT cells (GSM3130725 and GSM3130728) was performed using single-end mode implemented in Trimmomatic package (v. 0.36) using recommended settings and a minimum read length after trimming −20bp. Trimmed sequences were mapped to primary genome assembly GRCh38 using Bowtie2 mapper (v. 2.2.4). Sorted alignments were marked for PCR duplicates and used for downstream analyses. To prevent problems due to repetitive sequences that are collapsed in the current genome reference (and hence showing extremely high coverage by NGS reads), we first divided the genome into bins of 10 kb and determined the median number of reads mapping to a bin (primary alignments with a minimum mapping quality of 20, PCR duplicates excluded). All reads that mapped to (part of) a bin with either less than a tenth or more than 10 times the median number of reads per bin, were excluded from further analysis. Blacklists for each OK-seq dataset with these bins and the number of reads mapping there can be found in our code repository.

#### Correlating the OK-seq profiles of cells with and without replication stress

For each mapped OK-seq read (mapq >= 20, duplicates and secondary alignments were ignored) we determined the 5’ start location of the Okazaki fragment that this read must have originated from. We assumed that each fragment had a length of 150 bp. We used GSM3130725 as a reference to divide the autosomes into bins containing 10,000 Okazaki fragment 5’ starts. For each bin we determined the percentage of reads mapped to the forward strand in GSM3130725 and GSM3130728. Bins that contained less than 5000 fragment 5’ starts in GSM3130728 were excluded. The correlation between the percentage of fragments mapped to the forward strand in both samples was calculated using a Spearman rank test.

#### Identification of replication initiation zones using OK-seq data

Okazaki fragments come from the forward strand in the leftward moving fork of the origin, while they come from the reverse strand in the rightward moving fork. Replication initiation zones were identified by the detection of these strand usage switches using the script detect_strandswitches.pl. For each mapped read (mapq >= 20, duplicates and secondary alignments were ignored) we determined the 5’ start location and 3’ end location of the Okazaki fragment that this read must have originated from. We assumed that each fragment had a length of about 150 bp.

For each genomic region in between two consecutive Okazaki fragments we scored how many fragments mapped to the forward strand in the left flank (FL) and how many fragments mapped to the reverse strand in the right flank (RR), where the flank consisted of a hundred fragments (N =100 fragments per flank). An “origin score” was calculated as (FL + RR– N)/N, resulting in a score of zero for a random read distribution and a score of 1 for the perfect origin). Next, the regions were sorted by decreasing origin score. The region with the highest score was marked as a potential initiation zone, and regions for which the origin score was based on at least one of the same fragments as the identified potential initiation zone were subsequently excluded to prevent multiple calls in the same initiation zone. This process was repeated until there were no regions left to assign. When a replication initiation zone was identified in between two overlapping Okazaki fragments, we marked the middle base pair of the overlap. Finally, the significance of every potential initiation zone was determined using a binomial test where the number of successes was defined as FL + RR and the number of trials was the preset number of flanking fragments times two (N*2), using the script get_significant_oris.R.

The resulting p-values were corrected for the number of performed tests using the Benjamini-Hochberg method. Exclusion of potential initiation zones with FDR >= 0.01 and regions with a negative origin score (indicating potential fork merger zones) resulted in a final list of 10,733 identified replication initiation zones. A list of the replication initiation zones can be found in our code repository.

As a control we ran our Perl script while randomly shuffling strand information. Zero replication initiation zones were identified, confirming that our origin detection method is specific and does not identify replication initiation zones in random background data. Only autosomal replication initiation zones (n=9,608) were included into downstream analysis.

### Calculation of replication fork directionality at replication initiation zones

To examine the extent of replication continuation for each separate replication initiation zone, we calculated the OK-seq signal at the detected autosomal replication initiation zones using the script map_seqdata_around_oris_sliding_windows.pl and visualized the results in a heatmap. The signal was computed per 1 kb bins around each replication initiation zone, as the percentage of unambiguously mapped OK-seq reads that correspond to reverse strand fragments. Reads were assigned to bins according to the mapping coordinate of the central point of the fragment they originated from (assuming each Okazaki fragment length of about 150 bp). Due to low read coverage per individual initiation zone, we performed local smoothing, averaging signal from 4 bins upstream to 4 bins downstream (9 bins in total). The heatmap was plotted with the R package pheatmap 1.0.12.

We also calculated the average OK-seq signal across initiation zones using the script map_seqdata_around_oris, and plotted it. The signal was again computed per 1 kb bins around each replication initiation zone, as the percentage of unambiguously mapped OK-seq reads that correspond to reverse strand fragments. Alternatively, for figure S1D we calculated the RFD per 1kb bin as RFD=(F − R)/(F +R), where F and R correspond to the number of mapped reads to the forward and reverse strand, respectively. No smoothing was applied here. The signal was averaged across initiation zones.

### Raw data processing of double-click-seq data

The quality of sequencing data and potential contaminations were evaluated by FastQC software (v. 0.11.5). Quality trimming was performed using paired-end mode implemented in Trimmomatic package (v. 0.33). Trimmed sequences were mapped to primary genome assembly GRCh38 using Bowtie2 mapper (v. 2.2.4). Sorted alignments were marked for PCR duplicates and used for downstream analyses. Data have been deposited to EBI ArrayExpress under accession number E-MTAB-8624.

#### Calculation of new histone distribution bias at replication initiation zones

We calculated the average double-click-seq signal across initiation zones using the script map_seqdata_around_oris, and plotted it. The signal was computed per 1 kb bins around each replication initiation zone, as the percentage of unambiguously mapped read pairs that correspond to forward strand fragments. The calculation of partition score (Figure S1C) was performed per 1kb bins as (F −R)/(F +R), where F and R correspond to the number of mapped readpairs to the forward and reverse strand, respectively. Fragments were assigned to bins according to the mapping coordinate of their central points. Read pairs with a mapping quality below 20, discordantly mapped read pairs and read pairs corresponding to fragments shorter than 145 bp apart (that might not originate from a nucleosome-bound fragment) were excluded. The signal was averaged across initiation zones. When combining results from two replicates, averaging was first applied across initiation zones and then across replicates.

We also visualized the double-click-seq signal for selected individual replicates from untreated cells (replicate 1) and in cells treated with ATR inhibitor and hydroxyurea (replicate 2) around each separate replication initiation zone in heatmaps that can be seen in figures 1F and 3D. The script map_seqdata_around_oris_sliding_windows.pl was used for the calculations. The signal was again computed per 1 kb bins around each replication initiation zone, as the percentage of unambiguously mapped read pairs that correspond to forward strand fragments. Due to low read coverage per individual initiation zone, we performed local smoothing, averaging signal from 4 bins upstream to 4 bins downstream (9 bins in total). The result was visualized with the R package pheatmap 1.0.12.

#### Determining confidence intervals

Since initiation zones differ in terms of their constitutiveness and hence expected strand bias, we used bootstrap analysis for determining confidence intervals. For each replicate 1000 bootstrap replicates were performed to assess the robustness of observed strand bias. We used the range between the 2.5th and the 97.5th percentiles to show 95% confidence intervals on the plots. Scripts for this analysis can be found in our code repository.

#### Comparing bias at replication initiation zones in GC-rich versus AT-rich regions

The detected autosomal replication initiation zones were categorized into quartiles based on the GC percentage within 10 kb upstream and downstream of replication initiation zones (approximate distance at which we observe apparent strand asymmetry). The four quartiles had a GC percentage of below 35.6% (Q1), 35.6-37.4% (Q2), 37.4-40.3% (Q3) and above 40.3% (Q4). For each group of replication initiation zones the average histone deposition bias and replication fork directionality were calculated as described in sections above, with the exception that bin size was increased from 1 kb to 4 kb to compensate for lower read depth of quantile data using the script map_seqdata_around_oris_gc.pl.

#### Comparing bias at replication initiation zones in early replicated versus late replicated regions

The detected autosomal replication initiation zones were categorized into quartiles based on the average 100-cell mid-S RT scores of three hRPE-1 replicates for each initiation zone as downloaded from GEO accession GSE108556. Coordinates were lifted to hg38 using the UCSC Lift-over Genome Annotations tool with default settings. The four quartiles had an average 100-cell mid-S RT score of below −0.52 (Q1), −0.52 to −0.20 (Q2), −0.20 to 0.20 (Q3) and above 0.20 (Q4). A higher score indicates earlier replication. For each group of replication initiation zones the average histone deposition bias and replication fork directionality were calculated as described in sections above, with the exception that bin size was increased from 1 kb to 4 kb to compensate for lower read depth of quantile data using the script map_seqdata_around_oris_gc.pl.

#### Comparing bias at replication initiation zones in transcriptionally active versus inactive regions

The detected autosomal replication initiation zones were grouped into active or inactive regions based on hRPE-1 gene expression as downloaded from GEO accession GSE89413 (mapped to primary genome assembly GRCh38 using STAR aligner (v. 2.7.3) with --outSAMmapqUnique 50), using the script 02a_cluster_expressed.pl. Genes with a minimal expression of 1 FPKM and clusters of neighboring genes with a minimum expression of 1 FPKM each and maximum intergenic distance of 10 kb were regarded as active regions. The remaining sequences were marked as inactive regions. A bed file with the active and inactive regions can be found in our code repository. For each group of replication initiation zones the average histone deposition bias and replication fork directionality signal were calculated as described in section “Calculation of histone bias at replication initiation zones” and section “Calculation of replication fork directionality at replication initiation zones” respectively.

#### Multiple regression analysis to predict the effect of several features on histone deposition around initiation zones

For each double-click-seq sample we fitted the model y = β + β_1_a + β_2_b + β_3_c, where y= the new histone deposition bias at an initiation zone, a = the GC-fraction within 10 kb upstream and downstream of the initiation zone, b= the average 100-cell mid-S RT scores of three hRPE-1 replicates, c= transcriptional status (active or inactive) of the initiation zone. The strand deposition bias of new histones was calculated as the percentage of reads mapped to the forward strand in the 500:5500 bp region to the right of each origin zone minus the percentage of fragments from the forward strand in the −5500:−500 bp region to the left of each origin zone, divided by two. This means that a positive new histone deposition bias of maximally 50 indicates a lagging strand bias and a negative new histone deposition bias of maximally −50 indicates a leading strand bias.

## Reagents and Computational Resources

| REAGENT or RESOURCE | SOURCE | IDENTIFIER |
| --- | --- | --- |
| Azidohomoalanine | Santa Cruz Biotechnology | Cat#sc-285947; CAS 942518-29-8 |
| Biotin-PEG4-alkyne | Sigma-Aldrich | 764213-10MG; CAS 1262681-31-1 |
| THPTA | Sigma-Aldrich | 762342-100MG; CAS 760952-88-3 |
| CuSO<sub>4</sub> | Sigma-Aldrich | 451657-10G; CAS 7758-98-7 |
| Sodium ascorbate | Sigma-Aldrich | A7631-25G; CAS 134-03-2 |
| Dynabeads™ MyOne™ Streptavidin C1 | Thermo Scientific | 65801D |
| Dynabeads™ MyOne™ Streptavidin T1 | Thermo Scientific | 65801D |
| Azide-PEG3-biotin conjugate | Sigma-Aldrich | 762024-10MG; CAS 875770-34-6 |
| PARP inhibitor (Olaparib) | Axon Medchem | Axon 1464; CAS 763113-22-0 |
| Hydroxyurea | Sigma-Aldrich | H8627-1G; CAS 127-07-1 |
| Curaxin (CBL0137) | Selleckchem | Catalog No.S8483; CAS 1197397-89-9 |
| ATR inhibitor (VE-821) | Axon Medchem | Axon 1893; CAS 1232410-49-9 |
| MNAse | New England Biolabs | Cat# M0247S |
| pSpCas9(BB)-2A-GFP | Dr. Feng Zhang | N/A |
| NEBNext® Ultra™ DNA library prep kit for Illumina® | New England Biolabs | Cat# E7370S |
| NEBNext® multiplex oligos for Illumina® | New England Biolabs | Cat# E7335S (Index Primers Set 1) or Cat# E7500S (Index Primer Set 2) |
| Sequence reads | EBI ArrayExpress | E-MTAB-8624 |
| Custom Pearl Scripts | (https://github.com/thamarlobo/histone_deposition_analysis.git) | N/A |

### Biological Resources

hTERT immortalized human retinal pigmented epithelial cells (hTERT RPE-1, ATCC ®, CRL-4000™) were grown in T175 flasks at 37 °C with 5 % CO_2_ with DMEM, high glucose, GlutaMax™, pyruvate (Gibco) supplemented with 10% fetal bovine serum and 1% penicillin-streptomycin (10,000 units penicillin and 10 mg streptomycin/mL, Gibco). Cells were passaged with Trypsin-EDTA at 70-80% confluency and used for a maximum of 20 passages. Cells were regularly tested for mycoplasma infection.

## Results

### Double-click-seq reveals that bias of new histones is skewed towards the lagging strand

We used immortalized human retinal pigment epithelial (hRPE-1) cells to study histone deposition onto replicated DNA. The hRPE-1 cell line has a well-characterized genome and serves as a near-diploid model of normal human cells. The initial step for development of the double-click-seq protocol was optimization of the conditions for co-translational incorporation of AHA as a replacement of methionine into newly synthesized methionine-containing proteins over the time of one replication cycle in hRPE-1. We aimed to find conditions with minimal effect on the average cell doubling time (19). Cells were cultured in conditions with various AHA and methionine concentrations and histones were extracted and analyzed with mass spectrometry in order to assess the levels of AHA incorporation. Conditions were identified in which cells were cultured in a medium containing a mixture of 95% AHA and 5% methionine for 20 hours to reach a 3% AHA incorporation in histone H4 (Figure 1B), which were subsequently used throughout the study. The 20 hour labelling period allowed us to track histone deposition across a full cell cycle. Western blot analysis showed that AHA was also incorporated into histones H2A/B and H3 (Figure 1C).

After AHA labeling of *de novo* synthesized proteins, nuclei were isolated to remove cytoplasmic AHA-labelled proteins. The nuclei were then exposed to an initial copper-catalyzed [3+2] cycloaddition reaction between the azide group of AHA and an alkyne linked to biotin (click), which enabled affinity enrichment of AHA-labelled histones in complex with DNA (i.e. nucleosomes) and other AHA-labelled proteins using streptavidin beads after MNAse digestion. Subsequently, several washing steps were performed to rid all DNA not bound to histones. The remaining nucleosomal DNA was then extracted from the enriched nascent nucleosomes by digestion of histones and streptavidin by proteinase K treatment, after which the resulting enriched DNA was ligated to forked adaptors. In a second click reaction, nascent DNA strands, labeled with EdU (7), were coupled to an azide-linked biotin, thus enabling a second streptavidin bead capture. Finally, the unlabeled parental single stranded DNA was eluted from the beads with an alkaline solution and amplified to construct a directional short read sequencing library. The library was then sequenced with next generation sequencing and the resulting paired-end data (summary of all libraries in Table S1) were processed to assign genome-wide deposition of new histones to either the forward or reverse strand around the center of replication initiation zones (termed ‘origin centers’ in all related figures). Autosomal replication initiation zones were mapped using a publicly available Okazaki-fragment sequencing (OK-seq) data set for hRPE-1 cells (20) (n=9,608; Figure 1D and S1A).

Double-click-seq revealed that the deposition bias of new histones was skewed towards the strand replicated by lagging strand machinery at mapped replication initiation zones (Figure 1E, red line and Figure 1F). This finding is in agreement with the bias observed in mouse embryonic stem cells (7). Yet, our experiments indicate a more pronounced partition score as a measure of histone deposition bias of around 0.05 (calculated as proportion of forward and reverse read counts, see Figure S1C), whereas Petryk *et al*. reported a partition score of approximately 0.008. As we observe an average replication fork directionality (RFD) of 0.42 (Figure S1D), where *Petryk et al*. report an average RFD of around 0.13, variability in replication initiation zone firing rate between the different cell types used for both experiments can only partially explain the six-fold less pronounced asymmetry observed through immunoprecipitation of distinctive PTMs (7). The remaining difference might be the result of signal dilution through the exchange of histone PTMs over time, while our method stably labels new histones. Nonetheless, with double-click-seq we observed a clear bias of new histones around replication initiation zones towards the strand replicated by lagging strand polymerases. It is worth noting that our estimate of deposition bias might be conservative since turnover times range from fast for histones H2A/H2B to slow for H4 (0.4% per hour) (6).

We hypothesized that the deposition bias of new histones towards the lagging strand is most likely the result of the relatively less efficient capture of parental histones by the lagging strand, thus leaving the lagging strand with more new histones compared to the leading strand. Consequently, interfering with lagging strand synthesis would result in a more pronounced deposition bias of new histones towards the lagging DNA strand. During lagging strand synthesis, the 5’ flap of Okazaki fragments is excised by the nuclease FEN1, upon which the fragments are ligated by DNA ligase I (LIG1) (21). Under untreated conditions, a fraction of Okazaki fragments escapes LIG1-mediated ligation and is processed by a poly(ADP-ribose) polymerase (PARP)-mediated back-up route (22). Thus, inhibition of PARP activity interferes with the completion of lagging strand synthesis. In line with our hypothesis, we observed that treatment with the PARP inhibitor olaparib (23) increased the deposition bias of new histones onto the DNA strand replicated by lagging strand machinery (Figure 1E, yellow line). This indicates that disturbance of lagging strand synthesis increases the deposition bias of new histones to the lagging strand compared to the untreated condition.

### Replication stress inverts the general asymmetric deposition of new histones

Given the observed increase in the deposition asymmetry of new histones towards the lagging strand upon interference with lagging strand synthesis, we wondered what the effect of more general replication stress on new histone deposition would be. Strikingly, HU-induced replication stress completely inverted the bias of new histones towards the DNA strand replicated by leading strand synthesis (Figure 2A). An inversion of the deposition bias of new histones to the leading strand was also found upon treatment with the DNA intercalator curaxin (Figure 2B), which induces replication stress by triggering nucleosome unfolding (24). We checked if changes in origin usage may underlie the inverted histone deposition bias, however, we found that the OK-seq profiles between the untreated samples and HU treatment were highly correlated (Spearman correlation 0.98). Therefore, we conclude that replication stress, such as induced by either HU or curaxin treatment, inverts the asymmetry in new histone deposition relative to the untreated samples. Presence or absence of HU may explain the apparent contradiction between the leading strand bias of new histones found by Yu *et al*. (8) and the lagging strand bias of new histones found by Petryk *et al*. (7).

**Figure 2.**
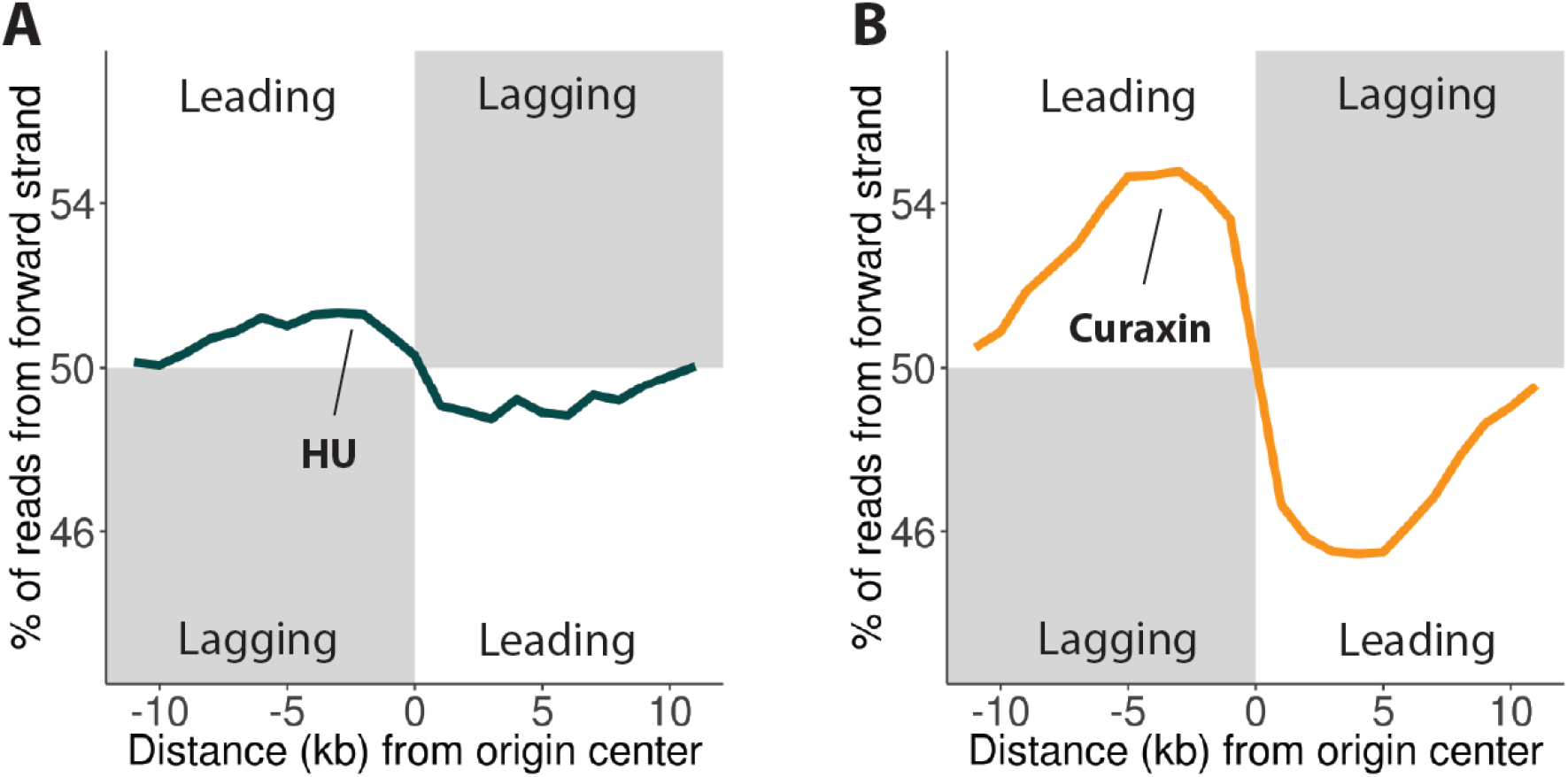
Replication stress inverts the inverts deposition asymmetry of new histones. (A) Average bias of new histone deposition at replication initiation zones during replication stress induced with hydroxyurea. (B) Average bias of new histone deposition at replication initiation zones during replication stress induced with curaxin. Separate plots for individual replicates including confidence intervals can be found in Supplementary Figure S2.

### Inhibition of DDR pathways also inverts new histone deposition asymmetry

As replication stress can lead to activation of the DDR, we speculated that the effect of replication stress on the deposition bias of new histones might be increased by inhibition of the DDR. The ataxia telangiectasia and Rad3-related (ATR) DDR pathway plays an essential role in suppressing replication stress (15). Additional DDR pathways involve the tumor suppressor p53 (16). We therefore investigated the involvement of the ATR- and p53-mediated DDR pathways in the altered histone deposition pattern during replication stress. Towards this aim, we employed p53-null hRPE-1 cells. Lack of p53 expression was confirmed by Western blot analysis (Figure 3A). In untreated p53-null cells, a deposition bias of new histones to the leading strand was observed (Figure 3B), indicating that loss of p53 already inverts the deposition bias of new histones seen in untreated hRPE-1 cells. The bias inversion increased further upon HU treatment of the p53-null cells (Figure 3B). Pharmacological inhibition of ATR in hRPE-1 cells also led to inversion of the deposition bias of new histones towards the leading strand. Again, HU treatment provided a stronger inversion of the deposition bias of new histones (Figure 3C). As an example, we included the heatmap of the deposition bias of new histones in cells treated with ATR inhibitor and HU, showing that histone bias inversion was observed consistently at the replication initiation zones that were identified (Figure 3D). We conclude that both replication stress and inhibition of DDR pathways through ATR or p53 inactivation disturb the asymmetry in the deposition of new histones such that the bias is inverted from the lagging to the leading strand.

**Figure 3.**
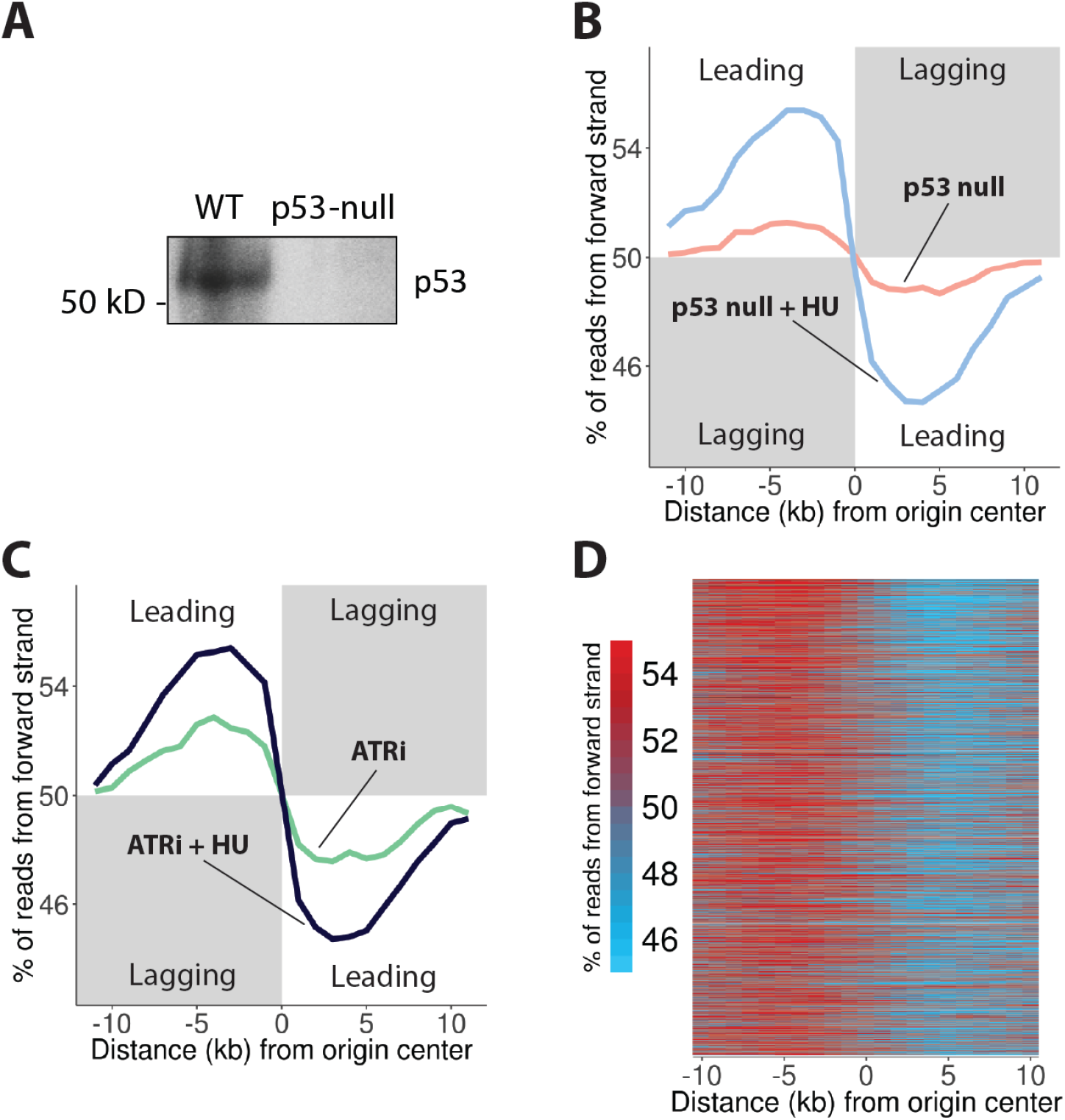
Inhibition of DDR pathways also inverts the deposition asymmetry of new histones. (A) Western blot showing knock-out of p53 in hRPE-1 cells with gene editing. Equal protein loading was ensured by measuring protein concentration with the Bradford assay. (B) Average bias of new histone deposition in p53-null hRPE-1 cells treated with hydroxyurea (light blue line) and without (pink line) at replication initiation zones. (C) Average bias of new histone deposition at replication initiation zones in hRPE-1 cells with either ATR inhibition (green line) or ATR inhibition and hydroxyurea treatment (dark blue line). (D) Heatmap of bias of new histone deposition at replication initiation zones in cells treated with ATR inhibitor and hydroxyurea. Separate bias plots of individual replicates including confidence intervals can be found in Supplementary Figure S2.

### Deposition bias of new histones is more pronounced in AT-rich regions

Finally, we investigated whether the histone deposition bias depends on common characteristics of genomic regions. We categorized replication initiation zones into actively transcribed (active) or not actively transcribed (inactive) (Figure 4C), early replicated or late replicated (Figure 4F), and AT-rich or GC-rich zones (Figure 4I). We then determined the average deposition bias of new histones around each category of initiation zones for both untreated hRPE-1 cells (‘untreated’, Figure 4A, D and G) and cells treated with ATR inhibitor and hydroxyurea (‘replication stress’, Figure 4B, E and H). We conclude that the deposition bias tends to be stronger in late replicated and AT-rich zones. Slight differences between replication initiation zones in actively transcribed versus not actively transcribed regions (Figure 4C) were observed too, but these differences were not large enough to account for the observed differences in histone deposition bias (Figure 4A and B).

**Figure 4.**
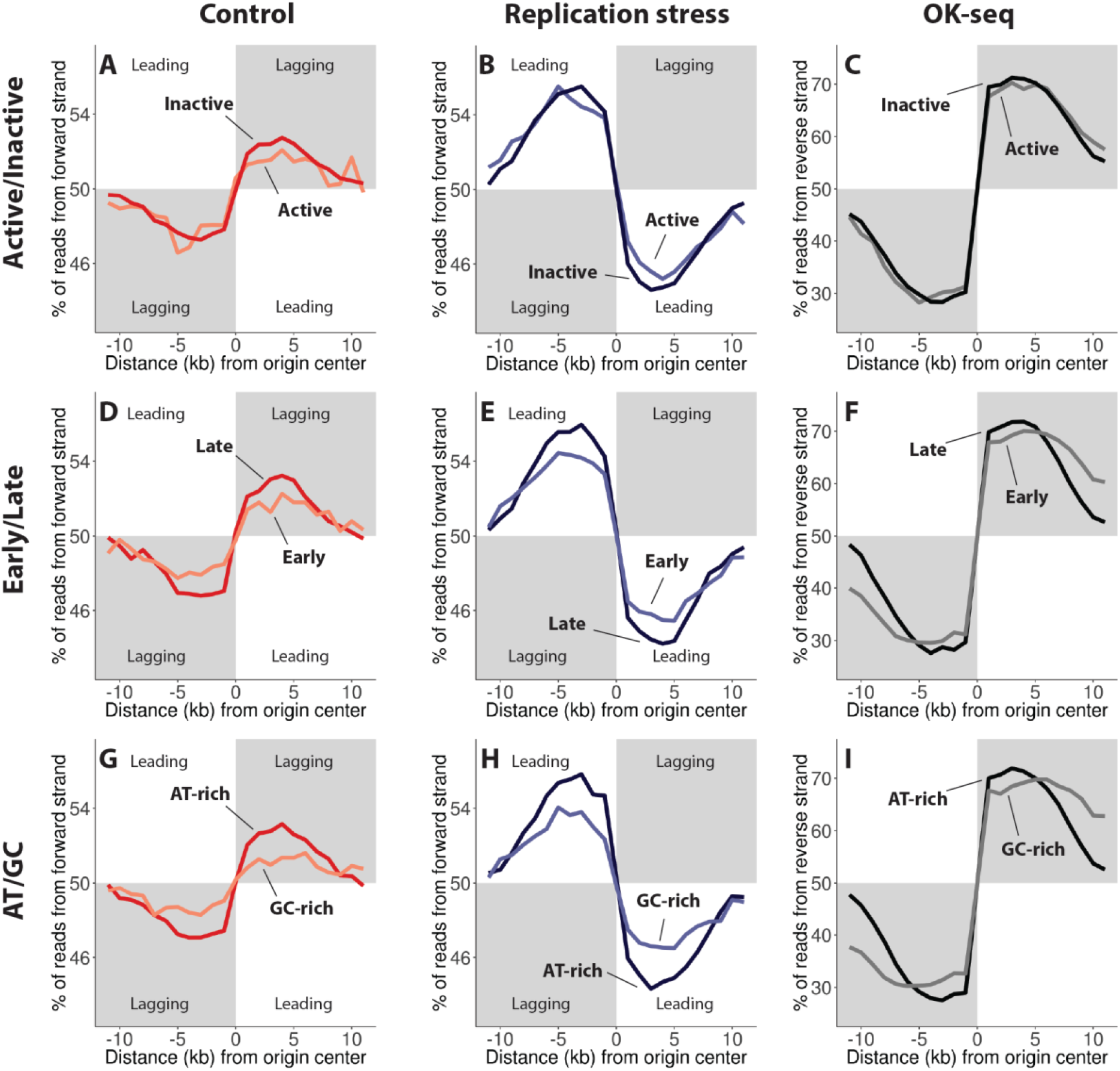
Deposition bias of new histones is more pronounced in AT-rich regions. Panels A, D and G show the average bias of new histone deposition at replication initiation zones in untreated conditions. Panels B, E and H show the average bias of new histone deposition at replication initiation zones with ATR inhibition and HU treatment. Panels C, F and I show the average replication fork directionality at replication initiation zones. For panels A, B and C the replication initiation zones were categorized into actively transcribed (red) and not actively transcribed (orange) regions. For panels D, E and F the zones were categorized into actively transcribed (light blue) and not actively transcribed (dark blue) regions. For panels G, H and I the zones were categorized into AT-rich (black) and GC-rich (grey) regions. Separate plots of individual replicates split into quartiles and including confidence intervals can be found in Supplementary Figure S3.

Genes that are actively transcribed are generally replicated in early S phase (25) and DNA regions replicated in early S phase generally have higher GC-content (26). In order to estimate the relative contribution of these genomic features to histone deposition asymmetry, we performed a multiple regression analysis taking all three factors into account (Table S2). GC-content proved to be the most consistently significant predictor of histone deposition bias. It was significant (uncorrected p-value < 0.01) for all used conditions, with the exception of the p53 knock-out cells. The GC-content beta values indeed signified a bias increasing effect in AT-rich zones across all conditions. Replication timing was a significant predictor for the bias in cells treated with ATR inhibitor, p53 knock-out cells and for one out of two replicates for cells treated with curaxin and cells treated with both ATR inhibitor and HU. Transcriptional status did not have a significant effect on the deposition bias of new histones (Table S2).

## Discussion

We developed the double-click-seq method to study the incorporation of new histones into nascent chromatin. We tracked *de novo* synthesized histones in cultured human cells and showed that they are enriched, under untreated conditions, in DNA replicated by the lagging strand machinery, especially in AT-rich regions. We found a more pronounced lagging strand bias in the deposition of new histones than reported previously (7) and we found this to be the case around all mapped replication initiation zones.

Most likely, redistribution of parental histones onto the DNA strand replicated by either leading or lagging strand polymerases favors the DNA strand that finishes replication first. The leading strand polymerase replicates DNA in a continuous fashion and directly follows unwinding of the parental DNA strands by the replicative helicase, such that the leading strand has the highest chance to capture a majority of the parental histones that are evicted ahead of the replication fork. The discontinuously replicated lagging strand by default would receive a majority of new histones to fill the gaps between re-deposited parental histones. In support of this model, we show that interfering with the completion of lagging strand synthesis with the PARP inhibitor olaparib increases the asymmetry in new histone deposition towards the lagging strand. Thus, with olaparib, the replicated lagging strand becomes less able to capture displaced parental histones and receives even more new histones than would normally be the case.

The asymmetry of new histone deposition towards the lagging strand underwent a striking inversion towards the leading strand upon treatment with the replication stress inducers HU and curaxin. Replication stress is known to induce uncoupling between the helicase and the leading strand polymerase, which leads to single-stranded DNA (ssDNA) that requires stabilization by binding of replication protein A (RPA) (27). Interestingly, the asymmetry in RPA occupancy, as an indirect measure of ssDNA, towards the lagging strand has been found to invert upon replication stress (27). Similarly, another study has shown that replication stress induces strand switching of the DNA clamp PCNA, such that more PCNA is found on the leading strand in stress conditions compared to untreated conditions (28). Both observations indicate that replication stress increases the proportion of ssDNA on the leading strand to a greater extent than on the lagging strand. The independently operating lagging strand polymerase is apparently better equipped to handle replication stress than its leading strand counterpart, perhaps in part due to repeated relocation of the lagging strand polymerase following completion of an Okazaki fragment (29). Therefore, when replication stressors target the replisome as a whole, the lagging strand polymerase is able to continue synthesis more effectively than the uncoupled leading strand polymerase, which is reflected in the bias inversions of RPA, PCNA and new histones. In essence, the DNA strand that is replicated by the lagging strand polymerase ‘leads’ during replication stress.

Accumulation of RPA on the leading strand following replication stress-induced helicase-polymerase uncoupling triggers a DDR that attempts to resolve the aberrant fork structure (13). Clearly, the DDR does not prevent inversion of new histone deposition asymmetry upon treatment with HU or curaxin. Previously, it has been demonstrated that chronic treatment of chicken DT40 cells with a low-dose of HU for 7 days stochastically induced a reduction in the expression of certain active genes, which was connected to a loss of the chromatin marks H3K4me3 and H3K9/14ac (11). It was postulated that the observed loss of histone marks was the result of uncoupling of histone recycling from DNA synthesis due to replication stress induced helicase-polymerase uncoupling. Our results provide more direct evidence for this hypothesis. Moreover, the results of Papadopoulou *et al*. illustrate that asymmetric deposition of new histones may accumulate over multiple cell divisions and cause epigenetic instability (11).

Additionally, we observed that functional DDR pathways are important in maintaining the characteristic pattern of histone deposition, since pharmacological inhibition of ATR and p53 knockout inverted the histone deposition pattern already without HU-induced replication stress. ATR is constitutively active in S phase to sense ongoing DNA replication and repress FOXM1 activity (30). After completion of S phase, a drop in ATR activity releases the brakes on FOXM1 activity and allows cells to enter mitosis. Apparently, inhibiting this intrinsic S/G_2_ checkpoint alters asymmetric histone deposition. Likewise, inactivation of *TP53* not only potentiated the observed bias inversion of new histones in HU-induced replication stress but also caused inversion by itself. This alludes to a constitutive function of p53 in maintaining replication fork integrity.

While OK-seq profiles for hRPE-1 cells treated with HU for 3 hours or without HU were strongly correlated, we did not determine the effect of 20 hour treatment with HU, the ATR inhibitor, olaparib nor p53 knock-out on OK-seq profiles. We can therefore not exclude the possibility that longer HU treatment and/or other treatments result in firing of more dormant origins, causing the asymmetry to seem weaker than it truly is. Additionally, some secondary cell divisions cannot be excluded because of the long labeling time. Such events might again weaken the asymmetry signal, but not enhance it nor change its direction.

Furthermore, we observe that histone deposition asymmetry is increased close to replication initiation zones located in AT-rich DNA. AT-rich DNA is less stable than GC-rich DNA due to a difference in hydrogen bonding (31). This difference in DNA stability leads to a higher rate of DNA unwinding in AT-rich compared to GC-rich regions, not taking the chromatic context into account (32). Such an increase in DNA unwinding rate may increase leading strand synthesis speed in untreated conditions and helicase-polymerase uncoupling during replication stress. In both cases this would lead to a greater difference in leading and lagging strand processivity in AT-rich compared to GC-rich regions and thus result in an increase in the asymmetry of new histone deposition.

In our hands, metabolic labelling of new histones with AHA is limited to approximately 3% of cellular histone H4 over the course of a 20 hours incubation period. All histone isoforms contain a methionine that can be replaced by AHA during *de novo* synthesis. Therefore, a labeling period that covers a full cell cycle leads to AHA incorporation in histones H3.3 and H2A/B, which have a replication-independent turnover (17). Histone H4 may also turnover independently from replication, but this seems mostly confined to centromeric regions (35). Regardless, replication independent turnover of histones could decrease the histone deposition asymmetry, but not enhance it nor change its direction. Also, we considered the possibility that the asymmetric distribution of newly synthesized proteins onto replicated DNA could reflect other proteins than histones, for example proteins that transiently associate with DNA during replication such as RPA and PCNA. We dismissed this possibility because 1) histones are by far the most abundant proteins associated with DNA at any time and 2) in our double-click-seq method we sequence DNA typically long after replication (by pulling down recently synthesized proteins and sequencing associated parental DNA template strands) and 3) we enrich for DNA fragments < 200 bp and exclude fragments with size < 145 bp from analysis. We also considered the fact that some old and newly synthesized histones will be exchanged after DNA replication before being pulled down in our method. Such post replication histone exchange events are expected to diminish the observed asymmetry in the distribution of new versus old histones over DNA replicated by leading versus lagging strand synthesis. Inversion of this asymmetry upon replication stress is furthermore difficult to reconcile with histone exchanges after DNA replication. For a more general analysis of strand-specific protein binding at replication forks we refer to the eSPAN method (28).

In conclusion, our observations indicate that the deposition of newly synthesized histones onto nascent chromatin is inherently biased towards the lagging strand, and that this bias increases upon PARP inhibition. Replication stress induced by HU or curaxin, genetic inactivation of p53 or pharmacological ATR kinase inhibition inverts the deposition bias of new histones towards the leading strand. These findings can be united by a model that includes differences in processivity of leading and lagging strand synthesis. Normally, polymerase ε on the leading strand is tightly coupled to the replicative helicase, whereas polymerase δ on the lagging strand is operating independently (33). As a result, re-deposition of parental histones is biased towards the faster completed leading strand and deposition of new histones is biased towards the lagging strand (Figure 5A). Helicase-polymerase uncoupling, due to a lack of polymerase ε processivity during replication stress, results in stretches of ssDNA on the leading strand (33, 34). The independently operating polymerase δ maintains processivity and therefore completes replication faster than polymerase ε (34). Together, this results in increased re-deposition of parental histones and decreased deposition of new histones on DNA replicated by polymerase δ, thus explaining the inversion of new histone deposition asymmetry towards the strand replicated by polymerase ε during replication stress (Figure 5B).

**Figure 5.**
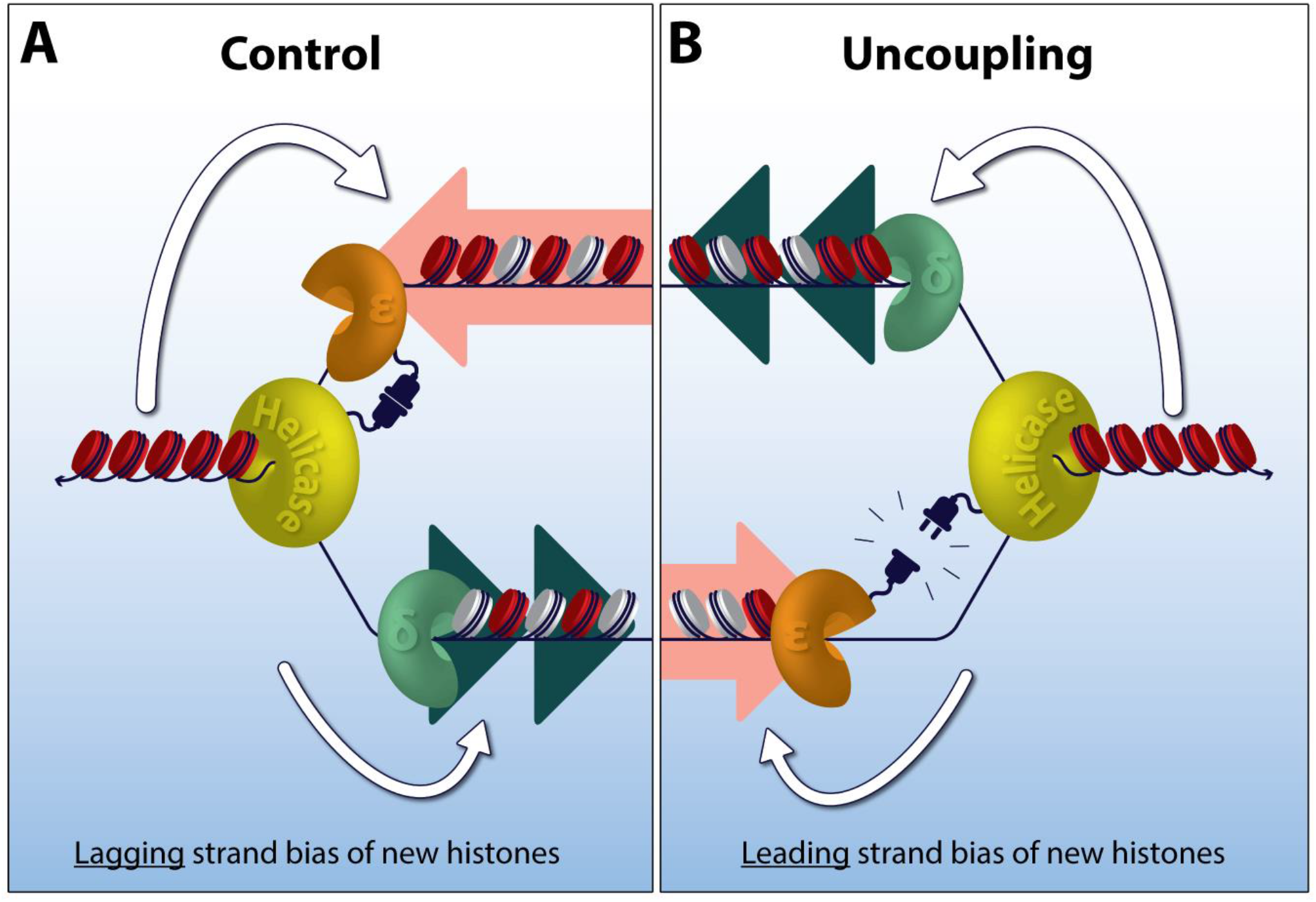
Helicase-polymerase uncoupling during replication stress inverts asymmetric histone deposition. (A) Model of lagging strand (green arrow heads) bias of new histones (white spheres) in untreated conditions following the leading strand (pink arrow) bias of old histones (red spheres), with polymerase ε (orange) coupled to the helicase (yellow). (B) Model of leading strand bias of new histone deposition following helicase uncoupling of polymerase ε. Helicase uncoupling results in lagging of leading strand synthesis. This results in a leading strand bias for new histones resulting from a biased re-deposition of old histones onto the faster completed lagging strand.

## Data Availability

The datasets generated during this study are available at EBI ArrayExpress under accession number E-MTAB-8624. Code generated during this study is available at GitHub; https://github.com/thamarlobo/histone_deposition_analysis.git

## Author Contributions

Conceptualization, P.M.L., F.J.D. and M.R.H.Z.; Investigation, methodology and validation, M.R.H.Z. and P.E.W. with help from D.C.J.S.; Data curation, formal analysis and software, T.J.L. and V.G.; Project administration, M.R.H.Z.; Funding acquisition, resources and supervision, F.J.D., V.G., D.C.J.S, P.M.L. and M.A.T.M.V.; Visualization, M.R.H.Z and T.J.L.; Writing – original draft, M.R.H.Z.; Writing – review & editing, all authors.

## Acknowledgments

We thank M. Groves for comments on the manuscript; and K. Hoekstra-Wakker, J. Beenen and N. Halsema for help with sequencing.

## References

1. Allis, C.D. and Jenuwein, T. (2016) The molecular hallmarks of epigenetic control. Nat. Rev. Genet., 17, 487.

2. Audergon, P.N.C.B., Catania, S., Kagansky, A., Tong, P., Shukla, M., Pidoux, A.L. and Allshire, R.C. (2015) Restricted epigenetic inheritance of H3K9 methylation. Science., 348, 132–135.

3. Ragunathan, K., Jih, G. and Moazed, D. (2015) Epigenetic inheritance uncoupled from sequence-specific recruitment. Science., 348.

4. Reverón-Gómez, N., González-Aguilera, C., Stewart-Morgan, K.R., Petryk, N., Flury, V., Graziano, S., Johansen, J.V., Jakobsen, J.S., Alabert, C. and Groth, A. (2018) Accurate Recycling of Parental Histones Reproduces the Histone Modification Landscape during DNA Replication. Mol. Cell, 72, 239-249.e5.

5. Escobar, T.M., Oksuz, O., Descostes, N., Bonasio, R. and Reinberg, D. (2019) Active and Repressed Chromatin Domains Exhibit Distinct Nucleosome Segregation During DNA Replication. Cell, 179, 953–963.

6. Alabert, C., Barth, T.K., Reverón-Gómez, N., Sidoli, S., Schmidt, A., Jensen, O., Imhof, A. and Groth, A. (2015) Two distinct modes for propagation of histone PTMs across the cell cycle. Genes Dev., 29, 585–590.

7. Petryk, N., Dalby, M., Wenger, A., Stromme, C.B., Strandsby, A., Andersson, R. and Groth, A. (2018) MCM2 promotes symmetric inheritance of modified histones during DNA replication. Science., 361, 1389– 1392.

8. Yu, C., Gan, H., Serra-Cardona, A., Zhang, L., Gan, S., Sharma, S., Johansson, E., Chabes, A., Xu, R.M. and Zhang, Z. (2018) A mechanism for preventing asymmetric histone segregation onto replicating DNA strands. Science., 361, 1386–1389.

9. Alexander, J.L. and Orr-Weaver, T.L. (2016) Replication fork instability and the consequences of fork collisions from rereplication. Genes Dev., 30, 2241–2252.

10. Chapman, T.R. and Kinsella, T.J. (2012) Ribonucleotide reductase inhibitors: A new look at an old target for radiosensitization. Front. Oncol., 1, 1–6.

11. Papadopoulou, C., Guilbaud, G., Schiavone, D. and Sale, J.E. (2015) Nucleotide Pool Depletion Induces G-Quadruplex-Dependent Perturbation of Gene Expression. Cell Rep., 13, 2491–2503.

12. Koç, A., Wheeler, L.J., Mathews, C.K. and Merrill, G.F. (2004) Hydroxyurea Arrests DNA Replication by a Mechanism that Preserves Basal dNTP Pools. J. Biol. Chem., 279, 223–230.

13. Byun, T.S., Pacek, M., Yee, M.C., Walter, J.C. and Cimprich, K.A. (2005) Functional uncoupling of MCM helicase and DNA polymerase activities activates the ATR-dependent checkpoint. Genes Dev., 19, 1040–1052.

14. Toledo, L.I., Altmeyer, M., Rask, M.B., Lukas, C., Larsen, D.H., Povlsen, L.K., Bekker-Jensen, S., Mailand, N., Bartek, J. and Lukas, J. (2013) ATR prohibits replication catastrophe by preventing global exhaustion of RPA. Cell, 155, 1088.

15. Zou, L. and Elledge, S.J. (2003) Sensing DNA damage through ATRIP recognition of RPA-ssDNA complexes. Science., 300, 1542–1548.

16. Klusmann, I., Rodewald, S., Müller, L., Friedrich, M., Wienken, M., Li, Y., Schulz-Heddergott, R. and Dobbelstein, M. (2016) p53 Activity Results in DNA Replication Fork Processivity. Cell Rep., 17, 1845–1857.

17. Deal, R.B., Henikoff, J.G. and Henikoff, S. (2010) Genome-Wide Kinetics of Nucleosome Turnover Determined by Metabolic Labeling of Histones. Science., 328, 1161–1164.

18. Ran, F.A., Hsu, P.D., Wright, J., Agarwala, V., Scott, D.A. and Zhang, F. (2013) Genome engineering using the CRISPR-Cas9 system. Nat. Protoc., 8, 2281–2308.

19. Arnaudo, A.M., Link, A.J. and Garcia, B.A. (2016) Bioorthogonal Chemistry for the Isolation and Study of Newly Synthesized Histones and Their Modifications. ACS Chem. Biol., 11, 782–791.

20. Chen, Y.H., Keegan, S., Kahli, M., Tonzi, P., Fenyö, D., Huang, T.T. and Smith, D.J. (2019) Transcription shapes DNA replication initiation and termination in human cells. Nat. Struct. Mol. Biol., 26, 67–77.

21. Balakrishnan, L. and Bambara, R.A. (2013) Okazaki fragment metabolism. Cold Spring Harb. Perspect. Biol., 5.

22. Hanzlikova, H., Kalasova, I., Demin, A.A., Pennicott, L.E., Cihlarova, Z. and Caldecott, K.W. (2018) The Importance of Poly(ADP-Ribose) Polymerase as a Sensor of Unligated Okazaki Fragments during DNA Replication. Mol. Cell, 71, 319-331.e3.

23. Menear, K.A., Adcock, C., Boulter, R., Cockcroft, X.L., Copsey, L., Cranston, A., Dillon, K.J., Drzewiecki, J., Garman, S., Gomez, S., et al. (2008) 4-[3-(4-Cyclopropanecarbonylpiperazine-1-carbonyl)-4-fluorobenzyl] - 2H-phthalazin-1-one: A novel bioavailable inhibitor of poly(ADP-ribose) polymerase-1. J. Med. Chem., 51, 6581–6591.

24. Safina, A., Cheney, P., Pal, M., Brodsky, L., Ivanov, A., Kirsanov, K., Lesovaya, E., Naberezhnov, D., Nesher, E., Koman, I., et al. (2017) FACT is a sensor of DNA torsional stress in eukaryotic cells. Nucleic Acids Res., 45, 1925–1945.

25. Hatton, K.S., Dhar, V., Brown, E.H., Iqbal, M.A., Stuart, S., Didamo, V.T. and Schildkraut, C.L. (1988) Replication program of active and inactive multigene families in mammalian cells. Mol. Cell. Biol., 10.1128/mcb.8.5.2149.

26. Woodfine, K., Fiegler, H., Beare, D.M., Collins, J.E., McCann, O.T., Young, B.D., Debernardi, S., Mott, R., Dunham, I. and Carter, N.P. (2004) Replication timing of the human genome. Hum. Mol. Genet., 13, 191–202.

27. Gan, H., Yu, C., Devbhandari, S., Sharma, S., Han, J., Chabes, A., Remus, D. and Zhang, Z. (2017) Checkpoint Kinase Rad53 Couples Leading-and Lagging-Strand DNA Synthesis under Replication Stress. Mol. Cell, 10.1016/j.molcel.2017.09.018.

28. Yu, C., Gan, H., Han, J., Zhou, Z.X., Jia, S., Chabes, A., Farrugia, G., Ordog, T. and Zhang, Z. (2014) Strand-Specific Analysis Shows Protein Binding at Replication Forks and PCNA Unloading from Lagging Strands when Forks Stall. Mol. Cell, 10.1016/j.molcel.2014.09.017.

29. Yu, C., Gan, H. and Zhang, Z. (2017) Both DNA Polymerases δ and ε Contact Active and Stalled Replication Forks Differently. Mol. Cell. Biol., 10.1128/mcb.00190-17.

30. Saldivar, J.C., Hamperl, S., Bocek, M.J., Chung, M., Bass, T.E., Cisneros-Soberanis, F., Samejima, K., Xie, L., Paulson, J.R., Earnshaw, W.C., et al. (2018) An intrinsic S/G2 checkpoint enforced by ATR. Science., 361, 806–810.

31. Yakovchuk, P., Protozanova, E. and Frank-Kamenetskii, M.D. (2006) Base-stacking and base-pairing contributions into thermal stability of the DNA double helix. Nucleic Acids Res., 34, 564–574.

32. Pandey, M. and Patel, S.S. (2014) Helicase and Polymerase Move Together Close to the Fork Junction and Copy DNA in One-Nucleotide Steps. Cell Rep., 6, 1129–1138.

33. Graham, J.E., Marians, K.J. and Kowalczykowski, S.C. (2017) Independent and Stochastic Action of DNA Polymerases in the Replisome. Cell, 169, 1201-1213.e17.

34. Georgescu, R.E., Yao, N., Indiani, C., Yurieva, O. and O’Donnell, M.E. (2014) Replisome mechanics: Lagging strand events that influence speed and processivity. Nucleic Acids Res., 42, 6497–6510.

35. Ahmad, K. and Henikoff, S. (2002) The histone variant H3.3 marks active chromatin by replication-independent nucleosome assembly. Mol. Cell, 9, 1191–200.

